# Chronic PPARγ Stimulation Shifts Amyloidosis to Higher Fibrillarity but Improves Cognition

**DOI:** 10.1101/2021.05.30.446348

**Authors:** Tanja Blume, Maximilian Deussing, Gloria Biechele, Finn Peters, Benedikt Zott, Claudio Schmidt, Nicolai Franzmeier, Karin Wind, Florian Eckenweber, Christian Sacher, Yuan Shi, Katharina Ochs, Gernot Kleinberger, Xianyuan Xiang, Carola Focke, Simon Lindner, Franz-Josef Gildehaus, Leonie Beyer, Barbara von Ungern-Sternberg, Peter Bartenstein, Karlheinz Baumann, Helmuth Adelsberger, Axel Rominger, Paul Cumming, Michael Willem, Mario M. Dorostkar, Jochen Herms, Matthias Brendel

**Affiliations:** DZNE – German Center for Neurodegenerative Diseases, Munich, Germany; Dept. of Nuclear Medicine, University Hospital of Munich, LMU Munich, Munich, Germany; Institute of Neuroscience, Technical University of Munich, Munich, Germany; Department of Diagnostic and Interventional Neuroradiology, Klinikum rechts der Isar, Technical University of Munich, Munich, Germany; Institute for Stroke and Dementia Research, University Hospital of Munich, LMU Munich, Munich, Germany; Metabolic Biochemistry, Biomedical Center (BMC), Faculty of Medicine, Ludwig-Maximilians-Universität München, Munich, Germany; ISAR Bioscience GmbH, Planegg, Germany; Roche Pharma Research and Early Development, Neuroscience Discovery, Roche, Innovation Center Basel, F. Hoffmann-La Roche Ltd., Basel, Switzerland; SyNergy, University of Munich, Munich, Germany; Dept. of Nuclear Medicine, Inselspital Bern, Bern, Switzerland; School of Psychology and Counselling, Queensland University of Technology, Brisbane, Australia; Center for Neuropathology and Prion Research, Ludwig-Maximilians-University of Munich, Munich, Germany

**Keywords:** pioglitazone, Aβ-PET, *App*^*NL-G-F*^ mice, PS2APP mice, microglia, Aβ-plaque composition

## Abstract

**Background:** We undertook longitudinal β-amyloid positron emission tomography (Aβ-PET) imaging as a translational tool for monitoring of chronic treatment with the peroxisome proliferator-activated receptor gamma (PPARγ) agonist pioglitazone in Aβ model mice. We thus tested the hypothesis this treatment would rescue from increases of the Aβ-PET signal while promoting spatial learning and preservation of synaptic density.

**Methods:** PS2APP mice (N=23; baseline age: 8 months) and *App*^*NL-G-F*^ mice (N=37; baseline age: 5 months) were investigated longitudinally for five months using Aβ-PET. Groups of mice were treated with pioglitazone or vehicle during the follow-up interval. We tested spatial memory performance and confirmed terminal PET findings by immunohistochemical and biochemistry analyses.

**Results:** Surprisingly, Aβ-PET and immunohistochemistry revealed a shift towards higher fibrillary composition of Aβ-plaques during upon chronic pioglitazone treatment. Nonetheless, synaptic density and spatial learning were improved in transgenic mice with pioglitazone treatment, in association with the increased plaque fibrillarity.

**Conclusion:** These translational data suggest that a shift towards higher plaque fibrillarity protects cognitive function and brain integrity. Increases in the Aβ-PET signal upon immunomodulatory treatments targeting Aβ aggregation can thus be protective.

## 1. Introduction

Alzheimer’s disease (AD) has become the most common cause of dementia, and is imposing a significant burden on health care systems of societies with aging populations (1). During the past few decades, research on AD pathogenesis led to the formulation of a model that accumulation of amyloid beta (Aβ)-plaques and neurofibrillary tangles, the histologically characterizing hallmarks of AD (2), triggers a cascade of neurodegenerative events, leading to disease progression (3). Additionally, novel emerging evidence indicates that neuroinflammation plays an important role in pathogenesis and progression of AD and many other neurodegenerative diseases (4; 5). In AD, activated microglial cells are able to bind and phagocytize soluble Aβ, and to some degree also the fibrillary Aβ aggregates, as part of the increased inflammatory response (4). However, others report that Aβ-recognition receptors on microglia downregulate during the progression of AD, such that microglial cells eventually undergo senescence, characterized by reduced phagocytosis of Aβ-aggregates (7). With time, the decreased microglial activity is permissive to expansion of fibrillar amyloidosis (8; 9) and a high proportion of dystrophic microglia were observed in human AD brain *post mortem* (11). These observations have led some to speculate that the microglial response is overwhelmed by the massive Aβ-deposition occurring in advanced AD, such that their chronic activation has a detrimental impact on disease progression (12; 7).

It might follow that treatment with anti-inflammatory drugs should alleviate AD progression. Pioglitazone is an anti-inflammatory insulin sensitizer widely used to treat hyperglycemia in type 2 diabetes via activation of peroxisome proliferator-activated receptor gamma (PPAR-γ). Treatment with pioglitazone enables microglial cells to undergo a phenotypic conversion from a pro-inflammatory towards an anti-inflammatory and neuroprotective phenotype (14; 15). Furthermore, activation of PPAR-γ in the brains of AD mice initiate a coupled metabolic cycle with the Liver X Receptor to increase brain apolipoprotein E levels, which promotes the ability of microglial cells to phagocyte and degrade both soluble and fibrillary Aβ (14; 15). However, another study showed that only low-dose PPAR-γ agonist treatment, but not the conventional doses, promotes an Aβ-clearing effect by increasing (LDL Receptor Related Protein 1 (LRP1) in human brain microvascular endothelial cells (HBMECs) (16). Despite this compelling preclinical evidence, a meta-analysis encompassing nine clinical studies did not compelling support a beneficial effect of PPAR-γ agonist treatment on cognition and memory in in patients with mild-to-moderate AD (18). Furthermore, a phase III trial of pioglitazone in patients with mild AD was discontinued due to lacking efficacy (19). It remains a conundrum why the translation of PPARγ stimulation into human AD failed, which calls for further investigation to uncover the basis of the seemingly false lead. Conceivably, the efficacy of pioglitazone may be confined to a specific stage of AD, or in cases distinguished by a particular biomarker.

Given this background, we hypothesized that Aβ-load and composition would determine the individual efficacy of PPARγ stimulation effect in the progression of AD mouse models. Therefore, we undertook serial small animal positron emission tomography (μPET) with the Aβ-tracer [^18^F]florbetaben (20–22) in two AD mouse models with distinct Aβ-plaque composition. The transgenic PS2APP-line develops dense fibrillary Aβ-plaques with late debit whereas the knock-In mouse model *App*^*NL-G-F*^ develops more diffuse oligomeric Aβ-plaques with early debut. Both strains of mice were treated with pioglitazone or vehicle for five months during the phase of main Aβ accumulation. We conducted behavioral assessments of spatial learning and confirmed longitudinal PET findings by immunohistochemical analysis and biochemical analysis, thus aiming to test the hypothesis that response to pioglitazone would depend on the type of Aβ-plaques formed in transgenic mice.

## 2. Methods and Materials

### Study design

Groups of PS2APP and *App*^*NL-G-F*^ mice were randomized to either treatment (PS2APP-PIO N=13; *App*^*NL-G-F*^-PIO N=14) or vehicle (PS2APP-VEH N=10; *App*^*NL-G-F*^-VEH N=23) groups at the age of 8 (PS2APP) and 5 (*App*^*NL-G-F*^) months. In PS2APP mice, the baseline [^18^F]florbetaben-PET scan (Aβ-PET) was performed at the age of eight months, followed by initiation of pioglitazone treatment or vehicle for a period of five months and a follow-up Aβ-PET scan at 13 months. In *App*^*NL-G-F*^ mice, the baseline Aβ-PET scan was performed at the age of five month, followed by initiation of pioglitazone treatment or vehicle, for a period of five months. Follow-up Aβ-PET scans were acquired at 7.5 months and ten months of age, which was the study termination in *App*^*NL-G-F*^ mice. For all mice, behavioral testing after the terminal PET scan was followed by immunohistochemical and biochemical analyses of randomized hemispheres. The TSPO-PET arm of the study and detailed analyses of neuroinflammation imaging are reported in a separate manuscript focusing on the predictive value of TSPO-PET for outcome of PPARγ-related immunomodulation (23). The sample size estimation of the in vivo PET study was based on previous experience and calculated by G*power (V3.1.9.2, Kiel, Germany), assuming a type I error α=0.05 and a power of 0.8 for group comparisons, a 10% drop-out rate per time-point (including TSPO-PET), and a treatment effect of 5% change in the PET signal (23). Shared datapoints between the study arms are indicated.

### Animals

PS2APP transgenic (24), *App*^*NL-G-F*^ APP knock-in (25) and wild-type C57Bl/6 mice were used in this investigation (for details see Supplement). All experiments were performed in compliance with the National Guidelines for Animal Protection, Germany, with approval of the local animal care committee of the Government of Oberbayern (Regierung Oberbayern) and overseen by a veterinarian. The experiments complied with the ARRIVE guidelines and were carried out in accordance with the U.K. Animals (Scientific Procedures) Act, 1986 and associated guidelines, EU Directive 2010/63/EU for animal experiments. Animals were housed in a temperature and humidity-controlled environment with a 12-h light–dark cycle, with free access to food (Ssniff) and water.

### Aβ-PET Acquisition and Reconstruction

[^18^F]florbetaben radiosynthesis was performed as previously described (22). This procedure yielded a radiochemical purity exceeding 98% and a specific activity of 80±20 GBq/μmol at the end of synthesis. Mice were anesthetized with isoflurane (1.5%, delivered via a mask at 3.5 L/min in oxygen) and received a bolus injection [^18^F]florbetaben 12±2 MBq in 150 μL of saline to a tail vein. Following placement in the tomograph (Siemens Inveon DPET), a single frame emission recording for the interval 30-60 min p.i., which was preceded by a 15-min transmission scan obtained using a rotating [^57^Co] point source. The image reconstruction procedure consisted of three-dimensional ordered subset expectation maximization (OSEM) with four iterations and twelve subsets followed by a maximum *a posteriori* (MAP) algorithm with 32 iterations. Scatter and attenuation correction were performed and a decay correction for [^18^F] was applied. With a zoom factor of 1.0 and a 128×128×159 matrix, a final voxel dimension of 0.78×0.78×0.80 mm was obtained.

### Small-Animal PET Data Analyses

Volumes of interest (VOIs) were defined on the MRI mouse atlas (26). A forebrain target VOI (15 mm^3^) was used for group comparisons and an additional hippocampal target VOI (8 mm³) served for correlation analysis with spatial learning. We calculated [^18^F]florbetaben standard-uptake-value ratios (SUVRs) using the established white matter (PS2APP; 67 mm^3^; pons, midbrain, hindbrain and parts of the subcortical white matter) and periaqueductal grey (*App*^*NL-G-F*^; 20 mm³) reference regions (27–29).

### Water Maze

Two different water maze tasks were applied due to changing facilities between the investigations of PS2APP and *App*^*NL-G-F*^ cohorts. We used a principal component analysis of the common read outs of each water maze task to generate a robust index for correlation analyses in individual mice (30). The principal component of the water maze test was extracted from three spatial learning read-outs (PS2APP: escape latency, distance, platform choice; *App*^*NL-G-F*^: escape latency, frequency to platform, time spent in platform quadrant). Thus, one quantitative index of water maze performance per mouse was generated for correlation with PET imaging readouts. The experimenter was blind to the phenotype of the animals.

#### Water Maze in PS2APP mice

PS2APP and age-matched wild-type mice were subjected to a modified Morris water maze task as described previously (31–34) yielding escape latency, distance to the correct platform and correct choice of the platform as read-outs.

#### Water Maze in App^NL-G-F^ mice

*App*^*NL-G-F*^ mice (treated and vehicle) and 14 age- and sex-matched wild-type mice (vehicle) underwent a classical Morris water maze test, which was performed according to a standard protocol with small adjustments (35) as previously described (29). Details are provided in the Supplement.

### Immunohistochemistry

Immunohistochemistry in brain regions corresponding to PET analyses was performed for fibrillary as well as oligomeric Aβ, microglia and synaptic density as previously published (36–38). We obtained immunofluorescence labelling of oligomeric Aβ using NAB228 (Thermo Fisher Scientific, USA) with a dilution of 1:500. For histological staining against fibrillar Aβ, we used methoxy-X04 (TOCRIS, Bristol, United Kingdom) at a dilution of 0.01 mg/ml in the same slice as for NAB228 staining. We obtained immunofluorescence labelling of microglia using an Iba-1 antibody (Wako, Richmond, USA) with a dilution of 1:200 co-stained with CD68 (BioRad, California, USA) with a dilution of 1:100. The synaptic density was measured using an anti-vesicular glutamate transporter 1 (VGLUT1) primary antibody (1:500, MerckMillipore). Quantification was calculated as area-%. Details are provided in the Supplement.

### Biochemical characterization of brain tissue

DEA (0,2% Diethylamine in 50 mM NaCl, pH 10) and RIPA lysates (20 mM Tris-HCl (pH 7.5), 150 mM NaCl, 1 mM Na2EDTA, 1% NP-40, 1% sodium deoxycholate, 2.5 mM sodium pyrophosphate) were prepared from brain hemispheres. The later was centrifuged at 14,000 g (60 min at 4°C) and the remaining pellet was homogenized in 70% formic acid (FA fraction). The FA fraction was neutralized with 20 x 1 M Tris-HCl buffer at pH 9.5 and used further diluted for Aβ analysis. Aβ contained in FA fractions was quantified by a sandwich immunoassay using the Meso Scale Aβ Triplex plates and Discovery SECTOR Imager 2400 as described previously (39). Samples were measured in triplicates.

### Statistics

The principal component of the water maze test was extracted using SPSS 26 statistics (IBM Deutschland GmbH, Ehningen, Germany). Prior to the PCA, the linear relationship of the data was tested by a correlation matrix and items with a correlation coefficient <0.3 were discarded. The Kaiser-Meyer-Olkin (KMO) measure and Bartlett’s test of sphericity were used to test for sampling adequacy and suitability for data reduction. Components with an Eigenvalue >1.0 were extracted and a varimax rotation was selected. Water maze results were also used as an endpoint in the dedicated manuscript on serial TSPO-PET in both cohorts (23). For immunohistochemistry quantifications GraphPad Prism (Graphpad Prism 7 Software, USA) was used. All analyses were performed by an operator blinded to the experimental conditions. Data were normally distributed according to Shapiro-Wilk or D’Agostino-Pearson test. One-way analysis of variance (ANOVA) including Bonferroni post hoc correction was used for group comparisons > 2 subgroups. For assessment of inter-group differences at single time points, Student’s t-test (unpaired, two-sided) was applied. All results are presented as mean ± SEM. P values <0.05 are defined as statistically significant.

## 3. Results

### Long-term pioglitazone treatment provokes a significant increase of the Aβ-PET signal in PS2APP mice

First, we analyzed serial changes of fibrillar amyloidosis under chronic pioglitazone treatment by [^18^F]florbetaben Aβ-PET in PS2APP mice and wild-type controls. Vehicle treated PS2APP mice showed an elevated Aβ-PET SUVR when compared to vehicle treated wild-type at eight (+20.4%, p<0.0001) and 13 months of age (+37.9%, p<0.0001). As expected, the Aβ-PET SUVR of wild-type mice did not change between eight and 13 months of age (0.831±0.003 vs. 0.827±0.008: p=0.645). Surprisingly, pioglitazone treatment provoked a stronger longitudinal increase in the Aβ-PET signal of PS2APP mice (+21.4%) when compared to vehicle treated PS2APP mice (+14.1%, p=0.002). At the follow-up time point, the Aβ-PET SUVR was significantly elevated when compared to untreated PS2APP mice (Fig. 1; 1.140±0.014 vs.1.187±0.011; p=0.0017). Pioglitazone treatment in wild-type mice provoked no changes of Aβ-PET SUVR compared to vehicle-treated wild-type mice at the follow-up time-point (0.827±0.008 vs. 0.823±0.005: p=0.496). Taken together, we found a significant increase in the Aβ-PET signal, which implied an increase in fibrillary Aβ-levels under pioglitazone treatment in PS2APP mice.

**Figure 1:**
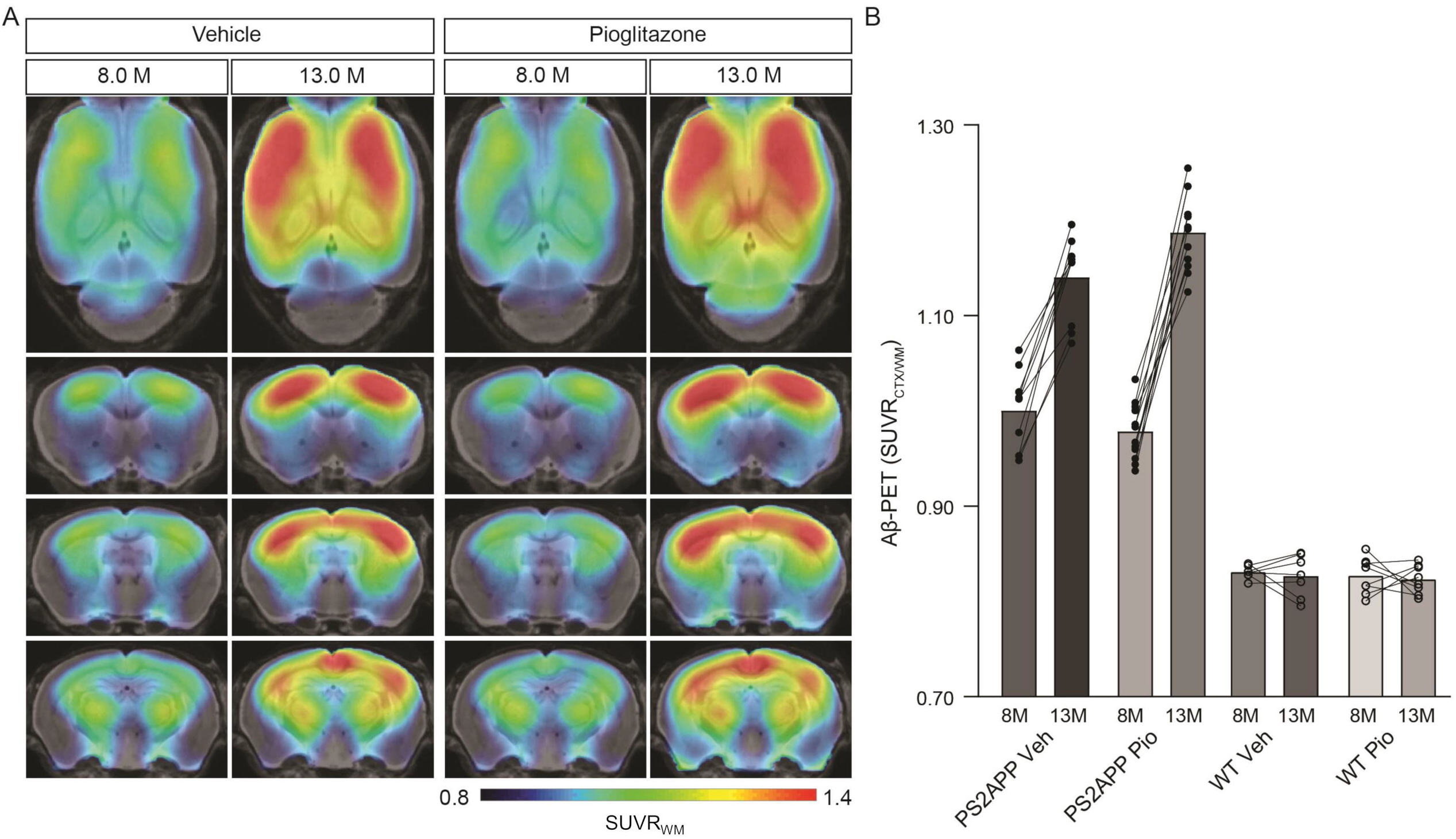
PPARγ stimulation in PS2APP mice provokes an increase in the Aβ-PET signal. A) Regional analysis of group-averaged standardized uptake value ratio (SUVR) images of the Aβ-PET radiotracer [^18^F]florbetaben in untreated and in pioglitazone-treated PS2APP mice aged eight and 13 months. Coronal and axial slices are projected upon a standard MRI template. B) Plots show cortical SUVR values of [^18^F]florbetaben in PS2APP and wild-type (WT) mice between eight and 13 months of age under vehicle (Veh) or pioglitazone (Pio) treatment. The Aβ-PET signal increased in PS2APP mice during aging, but the increase was more pronounced in pioglitazone treated mice (F_(1,12)_ = 12.9; p = 0.0017). In wild-type animals, no difference was observed between untreated and treated animals during aging (F_(1,13)_ = 0.490; p = 0.496). Data are presented as mean ± SEM. P values of Bonferroni *post hoc* test result from two-way ANOVA. N=10-13 PS2APP; N=7-8 WT.

### Aβ-PET detects a strong increase of the fibrillar Aβ-load in *App*^*NL-G-F*^ mice during chronic PPARγ stimulation

Next, we sought to validate our unexpected findings in PS2APP mice a mouse model with differing Aβ plaque composition, namely the *App*^*NL-G-F*^ mouse, which has limited fibrillarity due to endogenous expression of APP with three FAD mutations (25). Strikingly, the effect of pioglitazone treatment on the Aβ-PET signal was even stronger in *App*^*NL-G-F*^ mice than in PS2APP mice. There was a pronounced increase of the Aβ-PET signal during chronic pioglitazone treatment (+17.2%) compared to vehicle (+5.3%, p<0.0001). *App*^*NL-G-F*^ mice with pioglitazone treatment had a higher Aβ-PET SUVR at 7.5 (+4.6%, p=0.0071) and ten (+7.7%, p<0.0001) months of age when compared to vehicle-treated *App*^*NL-G-F*^ mice (Fig. 2). The baseline level of Aβ-PET SUVR was non-significantly lower in treated compared to untreated *App*^*NL-G-F*^ mice (0.878±0.010 vs. 0.906±0.006, p=0.1350). In both mouse models, the Aβ-signal increase after pioglitazone-treatment compared to baseline scans was pronounced in the frontotemporal cortex and hippocampal area (Figs. 1A & 2A). In summary, the pioglitazone treatment augmented the Aβ-PET signal increase in both mouse models; this unexpected result was more pronounced in the *App*^*NL-G-F*^ model, which expresses less fibrillary Aβ plaques.

**Figure 2:**
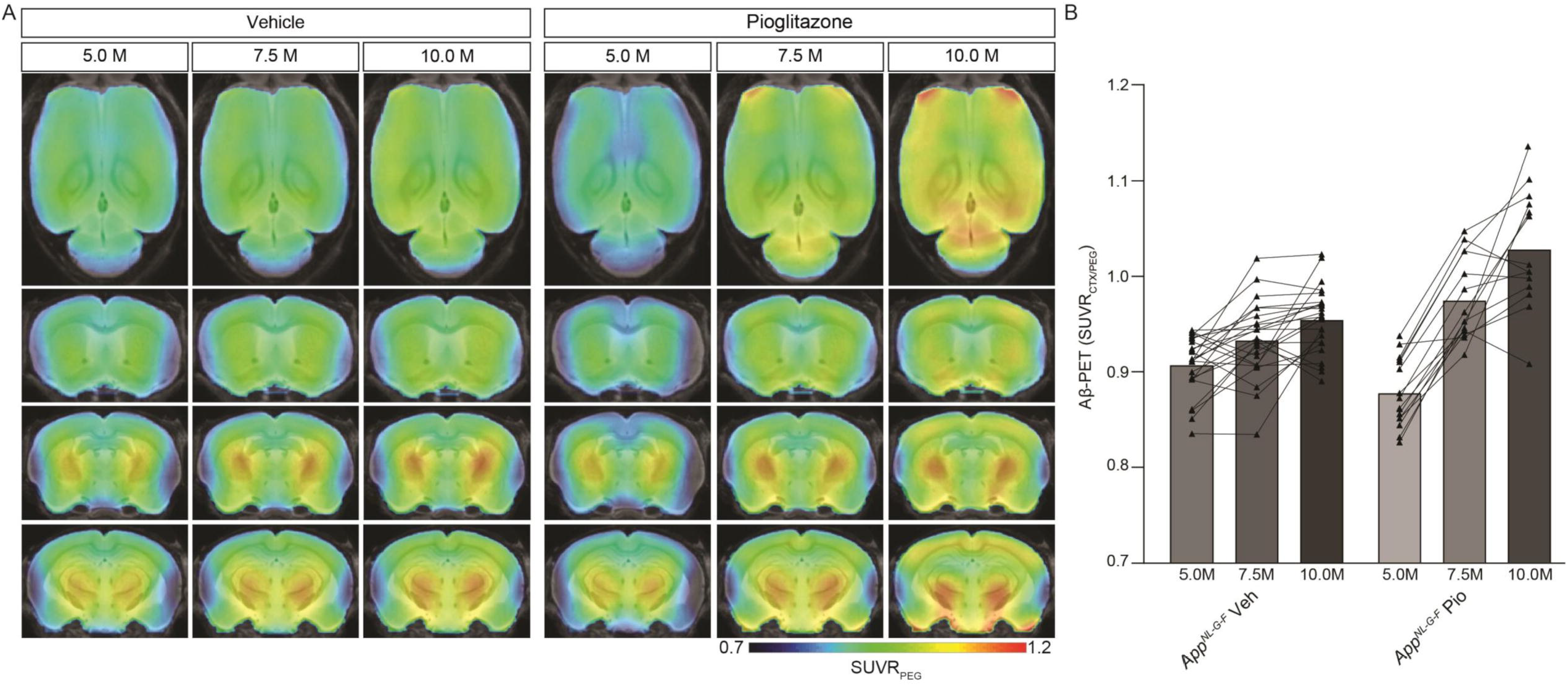
Distinct Aβ-PET signal increase upon PPARγ stimulation in *App*^*NL-G-F*^ mice with limited plaque fibrillarity and without overexpression of APP. A) Regional analysis of group-averaged standardized uptake value ratios (SUVR) of the Aβ-PET radiotracer [^18^F]florbetaben in untreated and in pioglitazone treated *App*^*NL-G-F*^ animals at the age of 5, 7.5 and 10 months. Coronal and axial slices are projected upon a standard MRI template. B) Plots show cortical SUVR of [^18^F]florbetaben in *App*^*NL-G-F*^ mice between the age of five and ten months under vehicle or pioglitazone treatment. Aβ-PET signal increased in untreated mice during age but the increase was more pronounced in pioglitazone treated *App*^*NL-G-F*^ mice (F_(2,70)_ = 20.12; p < 0.0001). Data are presented as mean ± SEM. P values of Bonferroni *post hoc* test result from two-way ANOVA. N=14-23.

### Pioglitazone triggers a shift towards increased Aβ-plaque fibrillarity in two distinct mouse models of amyloidosis

Given the unexpected *in vivo* findings, we set about to evaluate the molecular correlates of the potentiation of Aβ-PET signal during pioglitazone treatment in AD model mice. The (immuno)histochemical analysis showed that the observed increase of the Aβ-PET signal was predominantly explicable by a change in plaque composition rather than by a change in plaque density (Fig. 3). In both mouse models, the proportion of fibrillary Aβ stained with methoxy-X04 increased significantly under pioglitazone treatment compared to vehicle treated animals (PS2APP: 29.6±3.5% vs. 15.2±0.7%, p=0.0056, Fig. 3C; *App*^*NL-G-F*^: 9.1±1.6% vs. 4.4±0.4%, p=0.0001, Fig. 3D). Pioglitazone treatment had no significant effect on the proportion of oligomeric Aβ stained with NAB228 in PS2APP mice (PS2APP: 65.4±6.1% vs. 67.0±6.9%, p=0.865, Fig. 3C). In *App*^*NL-G-F*^ mice, however, the proportion of oligomeric Aβ decreased significantly in treated animals (*App*^*NL-G-F*^: 26.7±1.7% vs. 34.5±1.7%, p=0.0138, Fig. 3E). The effect size of pioglitazone treatment on plaque morphology was larger in *App*^*NL-G-F*^ mice than in PS2APP mice, which was reflected by a significantly increased overlay of methoxy-X04 and NAB228 positive plaques proportions in relation to untreated mice (PS2APP: 40.4±3.6% vs. 25.1±2.1%, p=0.0075, Fig. 3C; *App*^*NL-G-F*^: 35.0±3.4% vs. 12.9±1.3%, p=0.0005, Fig. 3E). We attribute this effect to the generally diffuse nature of the plaque composition of *App*^*NL-G-F*^ mice, which predominantly contain high oligomeric and low fibrillary fractions of Aβ (40) (compare Fig. 3A and Fig. 3B).

**Figure 3:**
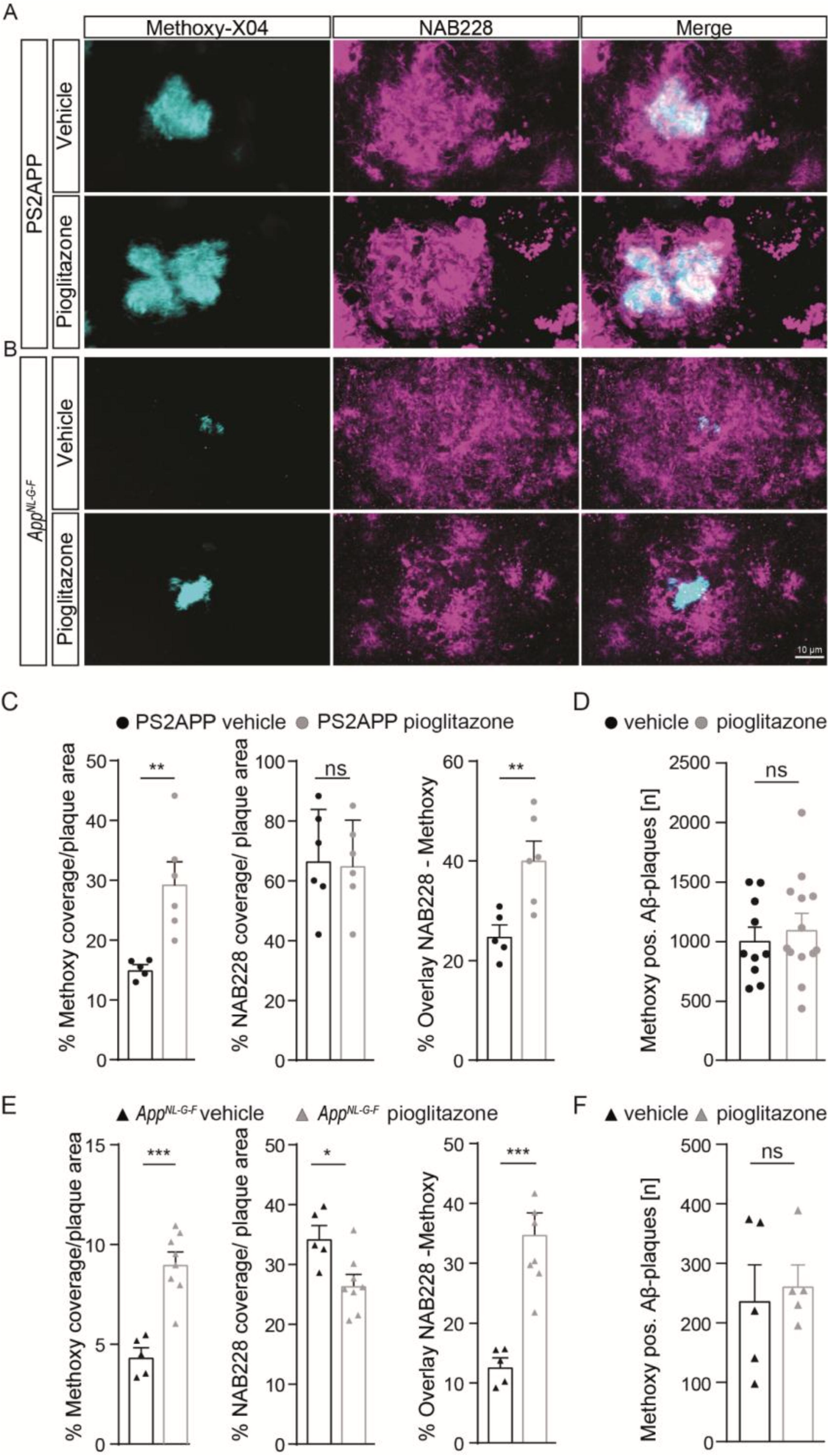
Pioglitazone treatment triggers a change in plaque composition in two different mouse models of amyloidosis. Staining of fibrillary Aβ (methoxy-X04, cyan) and oligomeric Aβ (NAB228, magenta) in vehicle and pioglitazone treated PS2APP mice A) and *App*^*NL-G-F*^ mice B). C) The plaque area covered by methoxy-X04 staining was significantly higher (t_(9)_ = 3.612; p = 0.0056), whereas the plaque area covered by NAB228 staining remained equal (t_(10)_ = 0.175; p = 0.865) in pioglitazone treated PS2APP mice. The overlay of NAB228 and methoxy staining increased under pioglitazone treatment (t_(9)_ = 3.432; p = 0.0075). D) The number of methoxy positive Aβ-plaques did not change under pioglitazone treatment in PS2APP-mice. E) In *App*^*NL-G-F*^ mice, methoxy coverage (t_(11)_ = 5.802; *p* = 0.0001), NAB228 coverage (t_(11)_ = 5.80; p = 0.0001), as well as the overlay of both stainings (t_(11)_ = 2.93; p = 0.0138), increased under pioglitazone treatment. F) In *App*^*NL-G-F*^ mice, the number of methoxy positive Aβ-plaques did not change under pioglitazone. Data are presented as mean ± SEM; n = 5-13 mice. Two-sample student’s *t*-test results: * p < 0.05; ** p < 0.01; *** p < 0.001.

The number of methoxy positive Aβ-plaques were similar between vehicle and pioglitazone treated groups for PS2APP (1016±107 vs. 1118±121, p=0.547, Fig. 3D) and *App*^*NL-G-F*^ mice (242±56 vs. 266±33, p=0.722, Fig. 3F). Notably there was no significant effect of chronic pioglitazone treatment on the different insoluble Aβ species (Aβ40, Aβ42) as well as on the level of the soluble Aβ42-isoform observed in either mouse model (Suppl. Fig. 1A). Taken together, our results indicate that the potentiated increase of the Aβ-PET signal upon pioglitazone treatment reflected a change in plaque composition from oligomeric to fibrillary Aβ-fractions.

### Microglial activation is reduced upon PPARγ stimulation in both AD mouse models

To confirm changes in the activation state of microglial cells, we performed Iba1 as well as CD68 immunohistochemical staining of activated microglia in both mouse models. We observed that pioglitazone treatment significantly decreased microglial activation in both mouse models (Fig. 4). In PS2APP mice, PPARγ stimulation provoked a one-third reduction of area coverage of Iba1-positive microglial cells (area: 9.1±0.6%) compared to untreated mice (14.0±0.5%, p=0.0003), and also a significant reduction of CD68-positive microglial cells area (7.6±0.4% vs. 9.9±0.3%, p=0.0018). In pioglitazone treated *App*^*NL-G-F*^ mice, the area reduction was less pronounced, but still significant for Iba1-positive microglial cells (9.4±0.2% vs. 10.6±0.2%, p=0.0015) and CD68-positive microglial cells (2.7±0.1% vs. 3.0±0.1%, p=0.0141) compared to untreated mice. Thus, we observed a consistent net reduction of activated microglial coverage in both models; the lesser effect in *App*^*NL-G-F*^ mice might indicate partial compensation by triggering of microglial activation due to increased fibrillary Aβ levels (40).

**Figure 4:**
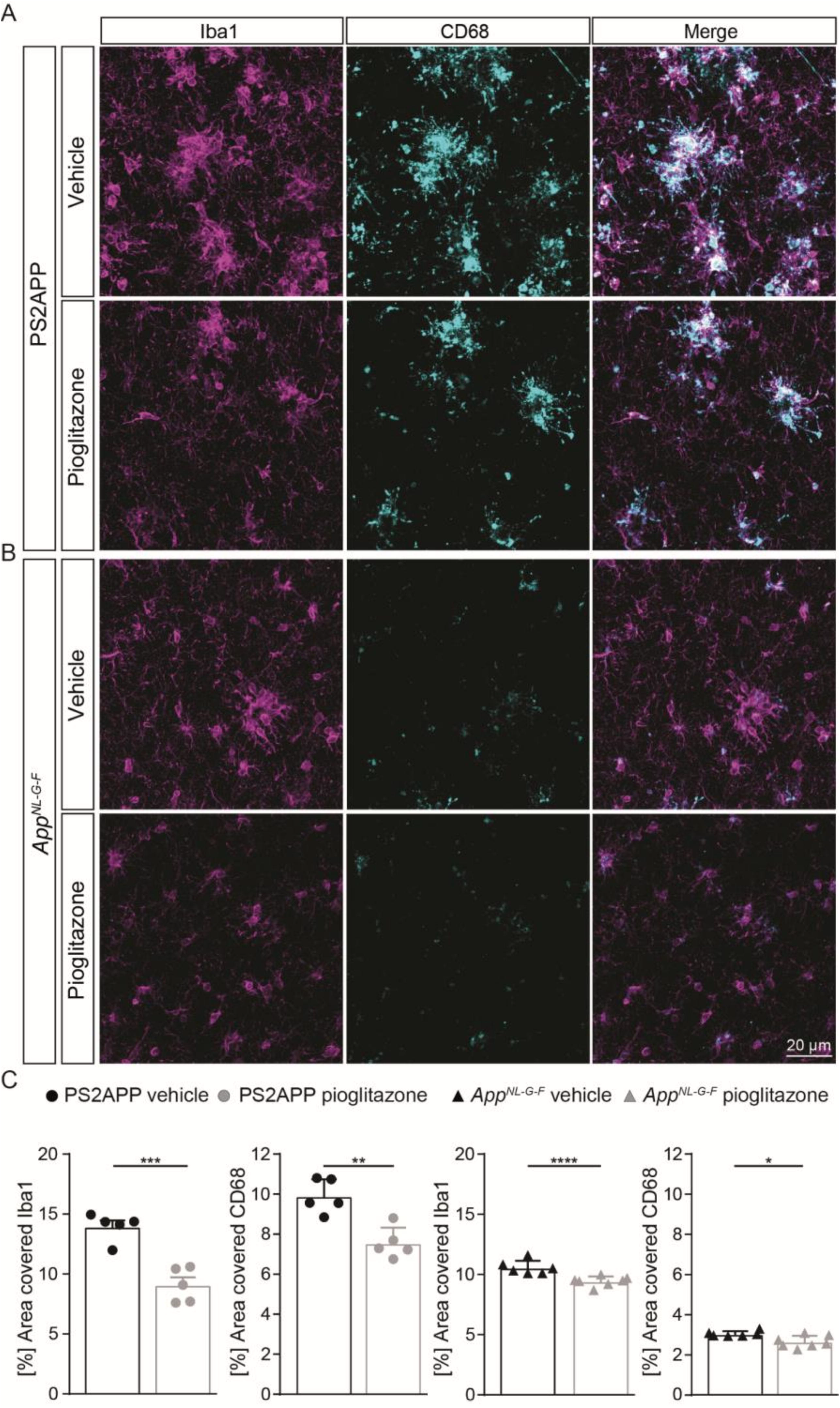
Pioglitazone treatment reduces microglial activation in both AD mouse models. Iba1-(magenta) as well as CD68-(cyan) positive microglial cells in PS2APP A) and *App*^*NL-G-F*^ mice B). C) The area of Iba1 positive microglial cells (t_(8)_ = 5.95; p = 0.0003) as well as CD68 positive microglial cells (t_(8)_ = 4.58; *p* = 0.0018) decreased in treated PS2APP mice. The same effect was observed in *App*^*NL-G-F*^ mice were the area covered by Iba1 positive (t_(11)_ = 4.21; *p* = 0.0015) as well as CD68 positive microglial cells (t_(11)_ = 2.91; p = 0.014) were significantly reduced in treated compared to untreated mice. Data are presented as mean ± SEM; n = 5-7 mice. Two-sample student’s *t*-test results: * p < 0.05; ** p < 0.01; *** p < 0.001.

### Cognitive function is improved by chronic pioglitazone treatment in association with an increasing Aβ-PET rate of change

Finally, we aimed to elucidate whether the observed longitudinal changes in the composition of Aβ-plaques affected synaptic density and hippocampus related cognitive performance. In PS2APP mice, treatment with pioglitazone resulted in a significant reduction of the water maze performance index compared to untreated mice during the probe trial (Fig. 5A; p=0.0155), whereas in wild-type animals there was no difference between treated and untreated animals (p>0.999). The water maze performance index of pioglitazone treated PS2APP mice correlated strongly with the rate of increase in Aβ-PET signal (Fig. 5C; R=0.686; p=0.0097). In *App*^*NL-G-F*^ mice, pioglitazone treatment did not result in a significant change of spatial learning performance (Fig. 5B; p>0.999). Accordingly, the water maze performance index and the rate of change in the Aβ-PET signal of pioglitazone treated *App*^*NL-G-F*^ mice did not correlate significantly (Fig. 5D; R=0.341; p=0.254). There was no significant association between the water maze performance index and the Aβ-PET rate of change in vehicle treated PS2APP or *App*^*NL-G-F*^ mice.

**Figure 5.**
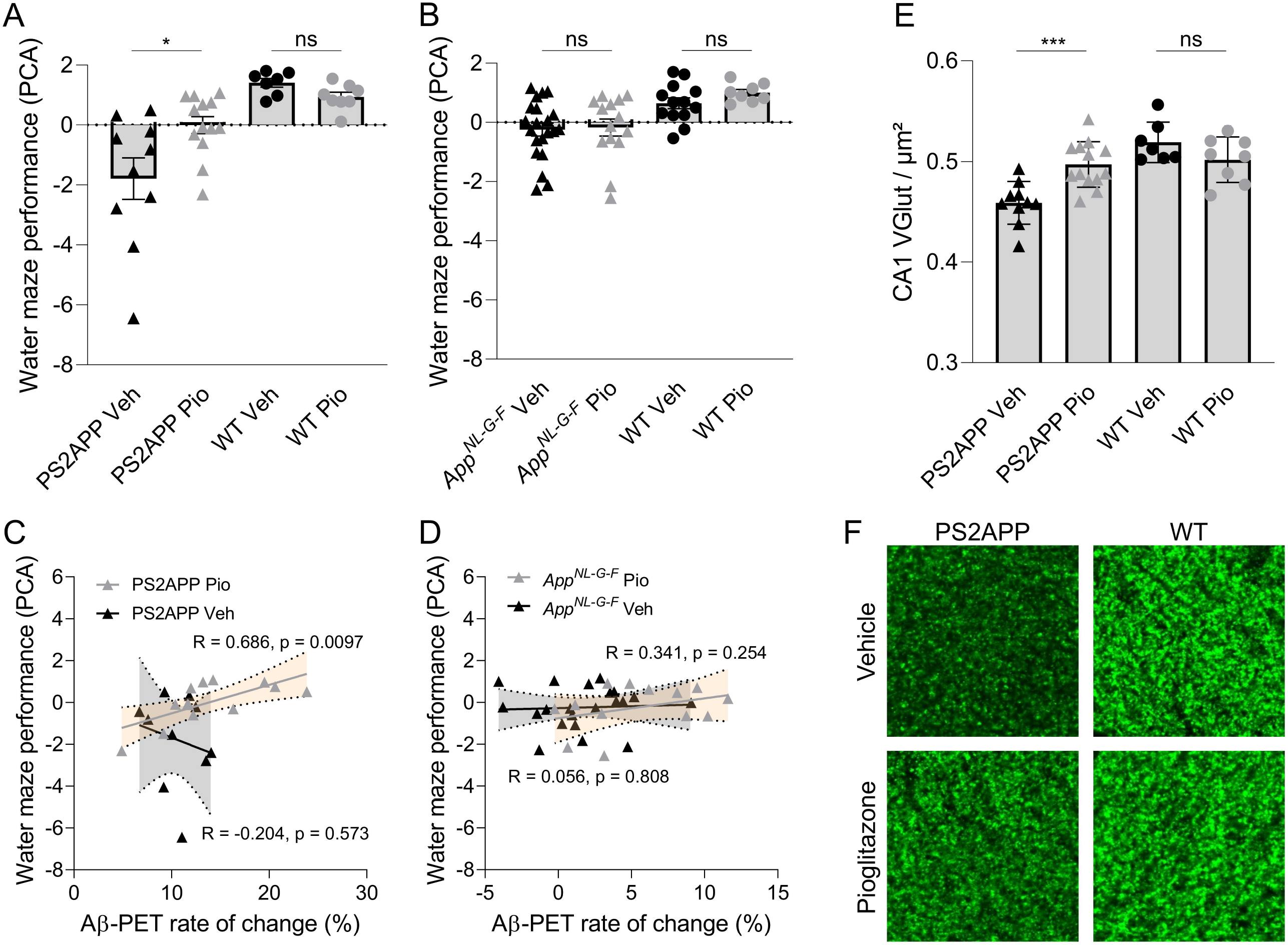
Improved spatial learning correlates with an increased Aβ-PET rate of change in PS2APP mice. A) One-way ANOVA revealed a significant difference of the water maze performance index between pioglitazone treated and untreated PS2APP and wild-type groups (F_(3,34)_ = 10.37; p < 0.0001; N=7-13). Group-wise comparisons revealed that pioglitazone treated PS2APP mice achieved a higher performance index in the water maze test compared to untreated PS2APP mice (p = 0.016), whereas wild-type animals showed no significant difference between treatment groups (p > 0.999). B) One-way ANOVA revealed a significant difference of the water maze performance index between pioglitazone treated and untreated *App*^*NL-G-F*^ and WT groups (F_(3,34)_ = 5.825; p = 0.0016). However, pioglitazone treated *App*^*NL-G-F*^ mice showed no difference in the water maze performance index when compared to untreated *App*^*NL-G-F*^ mice (p > 0.999) and wild-type animals again showed no significant difference between treatment groups (p > 0.999). Scatter plots show correlations between the Aβ-PET rate of change ([^18^F]florbetaben; ΔSUVR) during the treatment period and individual cognitive testing scores in C) PS2APP mice and in D) *App*^*NL-G-F*^ mice (R indicates Pearson’s coefficient of correlation) E) The decrease in synaptic density in the hippocampal CA1-region as assessed by VGLUT1 staining was ameliorated in treated PS2APP mice when compared to untreated mice (p = 0.0012), whereas no such treatment effect was seen in wild-type animals (p = 0.810; group effect: F_(3,34)_ = 12.03; p < 0.0001; N=7-13). F) VGLUT1 staining in the hippocampal CA1-region of representative untreated and treated PS2APP mice (left column) as well as of representative untreated and treated wild-type (WT) mice (right column). Statistics of group wise comparisons derive from one-way ANOVA with Bonferroni *hoc* correction: * p < 0.05; *** p < 0.005. Data are presented as mean ± SEM.

To explore the basis of water maze results in PS2APP mice at the molecular level, we performed staining of synaptic density in the hippocampus. Aβ-oligomers are the primary neurotoxic forms of Aβ, while Aβ-fibrils have less neurotoxicity (44–46). Thus, we hypothesized that pre-synaptic density in the hippocampal CA1-Area would be rescued upon pioglitazone-treatment. In wild-type mice we did not observe altered changed VGLUT1 density under pioglitazone treatment (Fig. 5E, F; 0.519±0.007 1/μm vs. 0.502±0.008 1/μm, p=0.810). In PS2APP mice, however, we found that pioglitazone treatment significantly rescued spine density in the CA1-region of the hippocampus compared to untreated animals (Fig. 5E, F; 0.497±0.006 1/μm vs. 0.459±0.007 1/μm, p=0.0012), supporting the hippocampal-dependent water maze results.

## 4. Discussion

To our knowledge, this is the first large-scale longitudinal PET study of cerebral Aβ-deposition in two distinct AD mouse models treated with the PPARγ agonist pioglitazone. We combined *in vivo* PET monitoring with behavioral testing and detailed immunohistochemical analysis. Our main finding was an unexpected potentiation in both mouse models of the increasing Aβ-PET signal during five months of pioglitazone treatment. This increase occurred despite an improvement of spatial learning and prevention of synaptic loss in the treated mice. Immunohistochemistry revealed a shift towards plaque composition of higher fibrillarity as the molecular correlate of the Aβ-PET signal, which was directly associated with improved cognitive performance in PS2APP mice.

Aβ-PET enables longitudinal *in vivo* detection of Aβ-plaques, which plays an important role in AD diagnosis, monitoring disease progression, and as an endpoint for therapeutic treatment effects (47). In our preceding observational and interventional studies, we validated in AD model mice the clinically established Aβ-PET tracer [^18^F]florbetaben relative to histologically defined indices Aβ deposition (3; 21). So far, an enhanced or increasing [^18^F]florbetaben-PET signal has been interpreted as an indicator of disease progression or treatment failure (48). Unexpectedly, we found that pioglitazone potentiated the increasing Aβ-PET signal in two mouse models compared to vehicle controls; in both cases, this increase was due to a shift of the plaque composition towards higher fibrillarity, and away from the more neurotoxic oliogomeric form. However, ELISA measurements of plaque associated fibrillary Aβ extracted with formic acid did not indicate a change in the Aβ species composition in brain. This suggests that Aβ-PET imaging and immunohistochemical analysis detect treatment effects on Aβ-plaque composition that do not arise from a shift in the levels of Aβ species, and which may thus evade detection in studies of CSF or plasma content (49).

Furthermore, our study provides evidence that rescued spatial learning deficits and prevented hippocampal synaptic loss can occur despite an increasing Aβ-PET signal upon immunomodulation. The combined results might sound contradictory, but according to the amyloid cascade hypothesis, Aβ-oligomers rather than Aβ-fibrils are the neurotoxic Aβ-forms (44; 50). Indeed, high concentrations of Aβ-oligomers isolated from brain of AD patients correlated significantly with the degree of cognitive impairment prior to death (51–53). Furthermore, Aβ-oligomers have been shown to disrupt long-term potentiation at synapses and provoke long-term depression (54–56). Thus, improved spatial learning and rescued synaptic density could reflect a therapeutically induced shift of Aβ to hypercondensed plaques, in keeping with observations of greater neuritic damage in association with more diffuse plaques (59; 60). Furthermore, strongly in line with our present data, a recent study argued that microglia promoted formation of dense-core plaques may play a protective role in AD (61).

The shift in plaque composition was more pronounced in *App*^*NL-G-F*^ mice than in the PS2APP model. Due to the expression of the Arctic mutation (25), the Aβ-deposits of the *App*^*NL-G-F*^ line consist predominantly of Aβ-oligomers (29; 40). However, we observed no improvement in cognition in the APP knock-in mouse line after pioglitazone treatment. We attribute the lacking improvement of spatial learning to the minor deterioration of this model in water maze assessment at ten months of age (64; 29). Our present observation stand in contrast with previous studies showing that PPAR-γ agonists reduced Aβ-plaque formation by increasing Aβ-clearance (15; 65; 14). However, those studies only performed endpoint analyses, in part after short-term treatment of nine days (14); the current work is the first to perform longitudinal *in vivo* monitoring of Aβ-deposition over a five-month chronic PPAR-γ treatment period. We note that the divergent results could also reflect the different markers used for immunohistochemistry compared to our present differentiated analysis of fibrillar and oligomeric Aβ components. As such, the decreased NAB228-positive plaque fraction in our treated *App*^*NL-G-F*^ mice fits to the earlier reported decrease of the 6E10-positive area in APPPS1 mice (14). We note that the biochemical source of the Aβ-PET signal is still a matter of controversy, since some studies found no impact of non-fibrillar plaque components (66) whereas others postulated a significant contribution of non-fibrillar Aβ to the Aβ-PET signal (67–69). Recently, we were able to show that non-fibrillar components of Aβ plaques indeed contribute to the net Aβ-PET signal (70). Therefore, increases in the [^18^F]florbetaben-PET signal must be precisely differentiated and interpreted with caution. Development of new PET tracers that selectively target oligomeric Aβ may realize a more precise discrimination of neurotoxic Aβ plaque manifestation (71; 72) and its impact on disease severity.

In line with previous pioglitazone studies (14; 15), we observed a decrease in microglial activity (23), thus confirming the immunomodulatory effect of the drug. Since earlier studies have shown that fibrillary Aβ-deposits activate microglial cells (40) which then migrate towards the fibrillar deposits (6), resulting in an increased number of activated microglial cells surrounding Aβ-plaques (8), the inactivation and migration effects could cancel each other out. Based on our findings in both AD models, we conclude that, by increasing plaque fibrillarity, the immunomodulatory effect of pioglitazone overweighs the potential triggering of activated microglia. Modulating microglial phenotype to restore their salutogenic effects may prove crucial in new therapeutic trials (74). In several preclinical and clinical trials, pioglitazone proved to be a promising immunomodulatory approach for treatment of AD, especially in patients with comorbid diabetes (75; 76). However, a large phase III trial of pioglitazone in patients with mild AD was discontinued due to lacking efficacy (19). Our data calls for monitoring of the effects of PPARγ agonists by Aβ-PET, which may help to stratify treatment responders based on their individual rates of Aβ plaque accumulation. Based on our results, we submit that personalized PPARγ agonist treatment might be effective when the patient has capacity to successfully shift toxic oligomeric Aβ towards fibrillar parts of the plaque.

## 5. Limitations

We note as a limitation that PPARγ receptor agonists represent a rather unspecific class of drugs since PPARγ is involved in various pathways in addition to peroxisome activation, notably including glucose metabolism and insulin sensitization [48]. Future studies should address if the observed effects on Aβ plaque composition are also present for more selective immunomodulation strategies such as NLRP3 regulators [49]. Two different water maze examinations were performed in the present study due a switch of the laboratory. Hence, although we calculated a similar water maze performance index by a PCA of the main read-outs of each examination, the obtained results and the sensitivity to detect spatial learning deficits are not comparable between both Aβ mouse models.

## 6. Conclusion

In conclusion, chronic pioglitazone treatment provoked a longitudinal Aβ-PET signal increase in transgenic and knock-in mice due to a shift towards hypercondensed fibrillar Aβ plaques. The increasing rate of Aβ-PET signal increase with time was accompanied by ameliorated cognitive performance and attenuated synaptic loss after pioglitazone treatment. It follows that increasing Aβ-PET signal need not always indicate a treatment failure, since it is the composition of Aβ plaques that determines their neurotoxiticy. In summary, our preclinical data indicate that a shift towards increasing fibrillar amyloidosis can be beneficial for the preservation of cognitive function and synaptic integrity.

## 7. Declarations

## Ethical Approval and Consent to participate

Not applicable

## Consent for publication

Not applicable

## Availability of data and materials

All source data are available from the corresponding author upon reasonable request.

## Author’s contributions

K.B., H.A., A.R., P.C., M.W., M.M.D., J.H. and M.B. conceived the study and analyzed the results. T.B., M.D. and M.B. wrote the manuscript with further input from all co-authors. M.D., G.B., C.Sch., K.W., F.E., C.Sa., and C.F. performed the small animal PET experiments and small animal PET data analyses. T.B., F.P., Y.S., K.O., G.K., X.X., M.M.D. and J.H. performed immunohistochemistry experiments, analyses, and interpretation. FJ.G. and S.L. performed PET tracer synthesis and analyses. N.F. analyzed and interpreted serial PET data and contributed to their analysis. G.B., B.Z., K.W., and H.A. performed spatial learning tests and interpretation. B.U.,K.B., and M.W. supplied the study with animal models and interpreted the dedicated results. All authors contributed with intellectual content.

## Acknowledgements

We thank Karin Bormann-Giglmaier and Rosel Oos for excellent technical assistance. Florbetaben precursor was provided by Piramal Imaging. We thank Takashi Saito and Takaomi C. Saido for providing the *App*^*NL-G-F*^ mice.

