## Supplement for "Chronic PPARγ Stimulation Shifts Amyloidosis to Higher Fibrillarity but Improves Cognition"

**Supplemental methods**

**Details on used mouse models**

The transgenic B6.PS2APP (line B6.152H) is homozygous for the human presenilin (PS) 2, N141I mutation and also the human amyloid precursor protein (APP) K670N, M671 L mutation. The APP and PS2 transgenes are driven by mouse Thy-1 and mouse prion promoters, respectively. This line had been created by coinjection of both transgenes into C57BL/6 zygotes. Homozygous B6.PS2APP (PS2APP) mice show the first appearance of plaques in the cortex and hippocampus at 5–6 months of age. The *App^NL-G-F^* mouse line carries a mutant APP gene encoding the humanized Aβ-sequence (G601R, F606Y, and R609H) with three pathogenic mutations, namely Swedish (KM595/596NL), Beyreuther/Iberian (I641F), and Arctic (E618G). Homozygous *App^NL-G-F^* mice show first appearance of plaques in the cortex and hippocampus at 3-4 months of age (25). Age-matched C57Bl/6 mice served as controls.

**Details on water maze experiments**

PS2APP: Mice had to distinguish between two visible platforms, one of which was weighted such that it would float when the mouse climbed on (correct choice), while the other would sink (wrong choice). The correct platform was always located at the same spot in the maze, while the wrong platform, as well as the site from which the mice were released into the maze, were varied in a pseudorandom fashion. Visual cues on the walls of the laboratory provided orientation. Trials were terminated if the mouse had failed to reach one of the platforms within 30 sec (error of omission). In this case, or in case of a wrong choice, the experimenter placed the mouse on the correct platform for 30 sec. After a three-day handling period, water maze training was performed on five consecutive days, with five trials per day, which were conducted 2-4 minutes apart. Memory performance was assessed by measuring the escape latency at each day of training and by the travelled distance at the last training day. For escape latency, we calculated the summed average time of all trials from the start point to attaining one of the platforms. On the sixth day, the right platform was placed in the opposite quadrant of the maze to confirm that the mice had used spatial cues rather than rule-based learning for navigation. Trials were filmed with a video camera and the swimming trace was extracted using custom-written LabView software (National Instruments).

*App^NL-G-F^*: The first day was used for acclimatization with the platform visible (five minutes per mouse). Then the mice underwent five training days in which each mouse had to perform four trials per day with the platform visible at the first training day, but submerged on all other training days. The test day consisted of only one trial with complete removal of the platform. The maximum trial length on all training and test days was set to a maximum of 70 seconds. The video tracking software EthoVision^®^ XT (Noldus) was used for analyses of escape latency, the platform visit frequency, and attendance in the platform quadrant at the probe trial.

**Details on immunohistochemistry**

Brains intended for immunohistochemistry were fixed by immersion in 4% paraformaldehyde at 4 °C for 15 hours. Two representative 50 µm thick slices per animal were then cut in the axial plane using a vibratome (VT 1000 S, Leica, Wetzlar, Germany). Free-floating sections were permeabilized with 2% Triton X-100 overnight and blocked with I-Block™ Protein-Based Blocking Reagent (Thermo Fischer Scientific, Waltham, USA). Slices were incubated for 24 hours at 4 °C. After three washes of ten min each in PBS, slices were incubated with an anti-guinea pig Alexa 488 secondary antibody (1:500, Life technologies) for four hours at room temperature. The unbound dye was removed in three washing steps with PBS, and the slices were then mounted on microscope slides with fluorescent mounting medium (Dako, Santa Clara, USA). Images were acquired with a LSM 780 confocal microscope (Zeiss, Oberkochen, Germany) equipped with a 40x/1.4 oil immersion objective. We acquired 3-dimensional 16-bit data stacks of 2048 x 2048 x 120 pixels from five different positions randomly selected in the frontal cortex, corresponding to the PET-VOIs, at alateral resolution of 0.17 um/pixel and an axial resolution of 0.4 μm/pixel for Iba1 and CD68 co-staining as well as for NAB228 and methoxy-X04 co-staining. Imaging of VGLUT1 staining was done in a scanned area of 42.43×42.43 µm^2^ with a lateral resolution of 0.083 µm in the CA1-region of the hippocampus. For each mouse, one VGLUT1 image per region was acquired. Images were further processed in Igor Pro to quantify the numbers of single puncta based on morphologic spot detection, as previously described (38). In brief, the sum of the unmixed second partial derivatives in the Cartesian coordinates x and y was calculated from an image and the sum was then thresholded at a 2×standard deviation of the pixel values to yield a binary image for determining and counting centers of mass for each punctum. Then, the numbers of pre-synapses were normalized to an adjusted image area by subtracting the areas occupied by cell bodies or blood vessels.

**Supplemental figures**

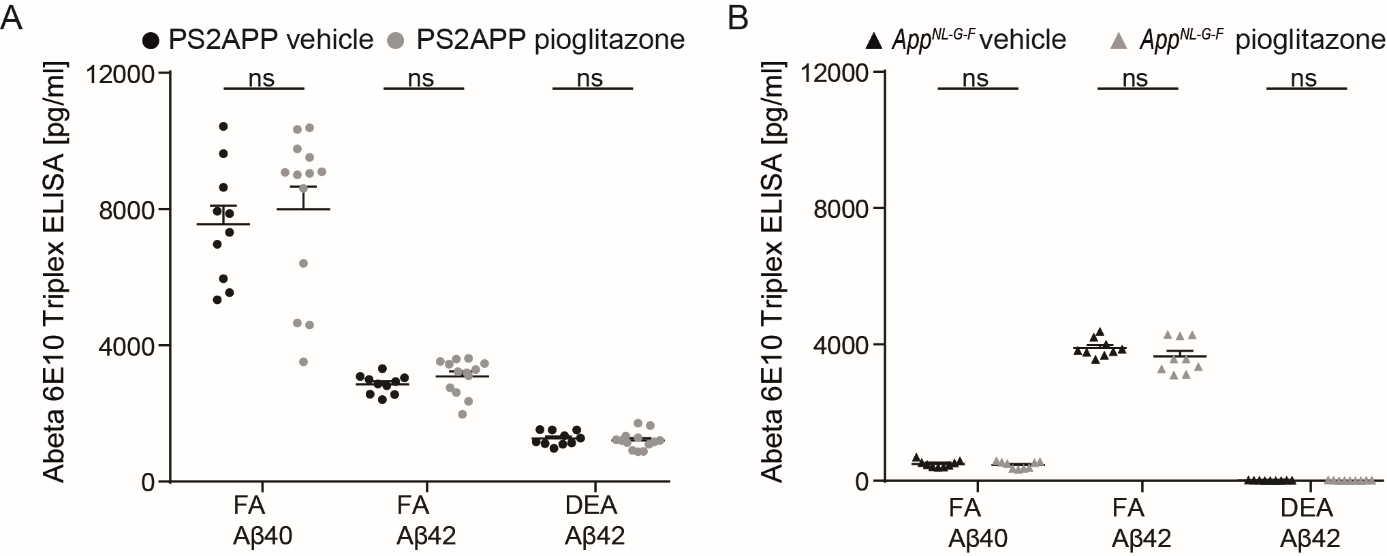

**Supplemental figure 1. Insoluble Aβ species as well as soluble Aβ42 did not change upon pioglitazone treatment in either mouse models.** A) ELISA-measurements showed a trend to changed insoluble Aβ-Isoform levels under pioglitazone treatment in PS2APP mice but did not reach significance for the total amounts of Aβ. In controls, 7560 (± 536) pg/ml Aβ40 was measured vs. 7999 (± 654) pg/ml Aβ40 in pioglitazone treated mice (p = 0.624, N=10) and nearly unchanged levels of 2856 (± 88) pg/ml Aβ42 in controls vs. 3089 (± 143) pg/ml Aβ42 in pioglitazone treated mice (p = 0.214; N=10). Levels of soluble Aβ42 did also not significantly change due to PPARγ-treatment. In controls 1265 (± 64) pg/ml soluble Aβ42 was measured vs. 1207 (± 71) pg/ml soluble Aβ42 in pioglitazone treated mice (p = 0.559, N=10; two-sample student’s t-test). B) No significant increase in the levels of insoluble Aβ-isoforms was observed due to PPARγ stimulation in *App^NL-G-F^* mice (control mice: 499 (± 33) pg/ml Aβ40 vs. pioglitazone treated mice: 466 (± 34) pg/ml Aβ40, p = 0.496, N=9 / control mice: 3902 (± 87) pg/ml Aβ42  vs. pioglitazone treated mice 3650 (± 167) pg/ml Aβ42, p = 0.198;  N=9; two-sample student’s t-test). No significant increase in the level of soluble Aβ42 was observed due to PPARγ stimulation in App^NL-G-F^ mice. In controls 12.1 (± 1.8) pg/ml soluble Aβ42 was measured vs. 11.4 (± 1.8) pg/ml soluble Aβ42 in pioglitazone treated mice (p = 0.789, N=9; two-sample student´s t-test).
